# Antifungal Resistance and Adhesin-Mediated Phenotypic Plasticity Among Genomically Diverse *Candida auris* Clinical Isolates

**DOI:** 10.64898/2026.08.26.747207

**Authors:** Tristan W. Wang, Tao Ma, Cindy Zhou, Rafael Gonzalez Martinez, Nicole Putnam, J. Kristie Johnson, Mary Ann Jabra-Rizk

## Abstract

*Candida auris* (currently *Candidozyma auris*) is an emerging fungal pathogen responsible for dramatic global increase in invasive candidiasis with high mortality. Most concerning, *C. auris* has a high propensity to colonize patients and persist and develop multidrug resistance to main classes of antifungals. In this study, we investigated the genetic and phenotypic diversity and resistance mechanisms of *C. auris* clinical isolates recovered from hospitalized infected patients. A total of 53 isolates from 38 unique patients were recovered from various clinical sources and evaluated for susceptibility to routine antifungal drugs. Whole genome sequencing (WGS) and single nucleotide polymorphism (SNP) analysis were performed to generate a phylogenetic network to infer population structure and identify mutations associated with drug resistance development. Isolates were also phenotypically evaluated for ability to form biofilms and aggregate, and cell wall adhesins gene expression studies were performed to provide mechanistic insights into *C. auris* phenotypic plasticity. Except for one clade III isolate, all isolates belonged to clade I and all were resistant to fluconazole with incidence of resistance to amphotericin B, echinocandins or both. Non-synonymous SNPs were found in genes associated with antifungal resistance including *ERG11, TAC1B, CDR1* and *FKS1*. Phenotypically, isolates varied in their ability to form biofilm and aggregate which correlated with expression of the Scf1 and Als4112 cell wall adhesins genes highlighting *C. auris* phenotypic plasticity in circulating clinical strains. These findings underscore the growing clinical threat posed by *C. auris* and reinforce the need for optimized surveillance and treatment strategies for controlling its spread.

## INTRODUCTION

*Candida auris* (currently *Candidozyma auris*) is an emerging nosocomial pathogen that has rapidly spread across continents causing outbreaks in healthcare facilities worldwide with an associated mortality rate of up to 60% (1-6). *C. auris* has 6 distinct clades with different genetic profiles and distinct geographic distributions and are reported to have differences in antimicrobial resistance, colonization and transmission rates (4, 6, 7). *C. auris* exhibits several concerning features including ability to persist in the hospital environment and to rapidly spread among patients (8-11). Due to its transmissibility, dramatic increase in cases and high levels of antifungal resistance, *C. auris* is categorized as an urgent threat and critical-priority pathogen by the Centers for Disease Control and Prevention and the World Health Organization (12).

In the United States, about 90% of *C. auris* isolates have been resistant to fluconazole, about 30% to amphotericin B and 0-8% to echinocandins with the South Asian clade I exhibiting the greatest percentage of resistant isolates (13). The echinocandins are currently the antifungal drugs of choice for treatment of *C. auris* infections, however, reports of resistance induced by antifungal therapy are on the rise, primarily associated with *FKS1* gene mutations (14-16). Similarly, resistance to amphotericin B was also reported to develop in patients on therapy although mechanisms of resistance are not well characterized (14, 17-19). Most concerning however, is the unique ability of *C. auris* to develop resistance to all classes of antifungals, severely limiting treatment options (1, 13, 20, 21).

Much like other *Candida* species, *C. auris* is able to form biofilms characterized by a dense consortia of adherent sessile cells embedded in an extracellular matrix that serve as a reservoir for the pathogen (22). Biofilm formation is an important virulence trait of fungal pathogens, as it enables adherence to biotic and abiotic surfaces and confers protection against antimicrobials (6). We previously classified *C. auris* isolates AR0382 and AR0387 belonging to the CDC AR-Bank as high-biofilm former and low-biofilm former, respectively (23). These isolates also differed in ability to aggregate. Cell aggregation is a unique phenotypic feature reported in some clinical *C. auris* isolates with aggregative and non-aggregative phenotypes exhibiting different biofilm forming abilities, drug susceptibility profiles and transcriptional changes (23, 24). The aggregative phenotype is predominantly in clade III strains and has been associated with skin colonization but there are also aggregative isolates in clade I and clade II (6, 25). To provide insight into this phenomenon, we recently analyzed the two phenotypes during biofilm growth in vitro and in vivo in a mouse model of catheter infection. Findings demonstrated significantly different transcriptional profiles and identified the cell wall adhesins Scf1 and Als4112 as key mediators of aggregation and biofilm formation (26).

Although the clinical relevance of aggregation is not fully understood, it has been reported that the *C. auris* phenotypes seem to have an association with virulence, with aggregative strains exhibiting reduced virulence in a mouse model of disseminated infection. Although the host environmental inducers remain unknown, *C. auris* was shown to rapidly evolve a multicellular aggregative morphology from single cells during systemic infection in a mouse model, with aggregates abundantly found in kidney tissue (27, 28). These findings suggest that aggregate morphology may be associated with increased fungal burden and tissue colonization. In fact, aggregative strains were also shown to resist clearance by the immune system, suggesting that aggregation might facilitate long-term persistence in the host (25).

More recently, we reported on a clinical case of *C. auris* multi-drug resistance development in a transplant patient receiving antifungal therapy and identified a mutation in the *FKS1* gene associated with echinocandin resistance (14). However, no mutations were found associated with amphotericin B resistance, indicating unknown genetic elements likely driving resistance. Intriguingly, although recovered from the same patient, the isolates displayed phenotypic variations in growth, aggregation and biofilm formation. In this study, we leveraged a collection of *C. auris* clinical isolates recovered from hospitalized infected patients and performed whole genome sequencing and SNP calling to generate a phylogenetic network to infer population structure. Further, we characterized the phenotypic diversity and genetic determinants of antifungal resistance, and importantly, we performed cell wall adhesins gene expression studies to provide mechanistic insights into *C. auris* phenotypic plasticity.

## MATERIALS AND METHODS

### *C. auris* clinical isolates

The study was approved by the University of Maryland, Baltimore Institutional Review Board (HP-00110276) and Institutional Biosafety Committee (IBC-00007835). Isolates were obtained from deidentified clinical samples recovered from patients attending the University of Maryland Medical System (11 hospitals) between April 2021 and June 2025. A total of 53 isolates were recovered from various clinical sources (Table 1). Isolates were obtained from 38 unique patients, 8 of which had multiple (2-5) isolates recovered. *C. auris* isolates were identified by matrix-assisted laser desorption/ionization time-of-flight mass spectrometry (MALDI-TOF MS) using the VITEK MS (bioMérieux). Isolates were stored in 20% glycerol stocks at -80°C and grown on yeast peptone dextrose agar (YPD) (Difco Laboratories) at 37°C. The AR0387 (B8441) reference strain was included in some studies as.

**Table 1.**
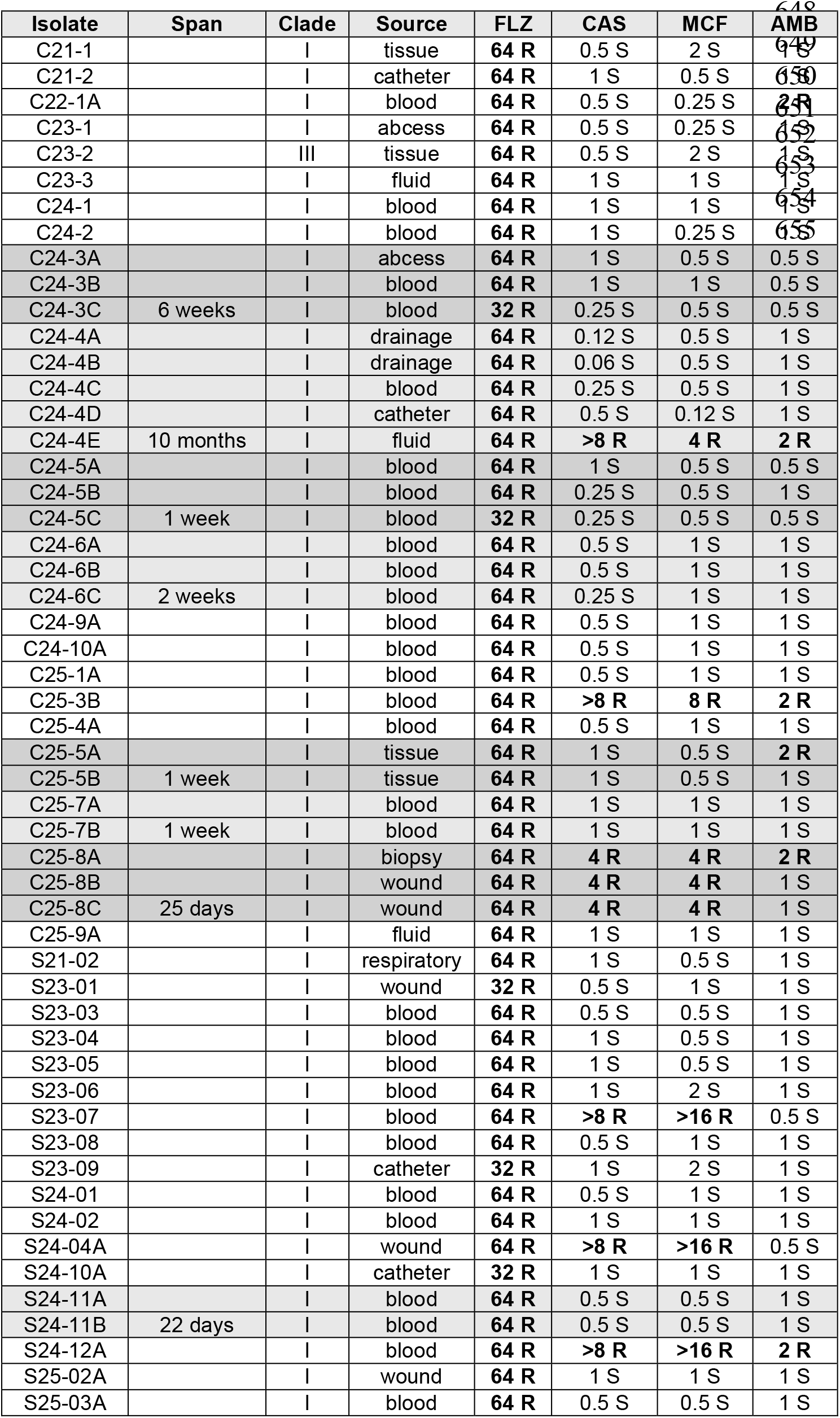
Characteristics of clinical isolates with antifungal MICs (µg/ml). Boxed isolates are recovered from same patient; span indicate time between first and last recovered isolates in days, weeks or months. MICs shown in bold exceed tentative CDC MIC (μg/ml) breakpoints for resistance: fluconazole (FLZ) ≥32; amphotericin B (AMB) ≥2; caspofungin (CAS) ≥2; micafungin (MCF) ≥4.

### Antifungal susceptibility testing

MICs for amphotericin B, fluconazole, caspofungin and micafungin were determined using the Clinical and Laboratory Standards Institute (CLSI) microdilution method (Supp M60). MICs were read visually after 24-48h incubation at 35°C and resistance determined based on MICs exceeding tentative CDC breakpoints for *C. auris (*https://www.cdc.gov/candidaauris/site.html; accessed June, 2026).

### Whole genome sequencing

Genomic DNA was extracted from overnight YPD broth cultures using the DNeasy Blood and Tissue extraction kit (Qiagen) according to manufacturer instructions with an added bead beating step. Sequencing libraries were generated with the NEBNext Ultra II DNA Library Prep Kit and sequenced on a 25B flowcell on the Illumina NovaSeq X Plus instrument using a 2×150 paired-end (PE) configuration, targeting ∼1 Gb/sample. bcl files generated by the sequencer were converted into fastq files and de-multiplexed using Illumina’s bcl2fastq 2.20 software.

### Phylogenetic analysis and variant calling

Trimmomatic v0.40 was used to remove sequence adapters and poor-quality reads. BWA-MEM v0.7.19 (29) was used to align trimmed reads to the clade I reference genome B8441 (GenBank accession GCA_002759435.3). Mean sequencing depth was greater than 55x for all isolates. Paired-end reads for the clade III reference genome B11221 (Sequence Read Archive run accession SRR3883453) were processed identically to the clinical isolates. Variants were called from the aligned reads using bcftools mpileup with minimum mapping quality and minimum base quality of 20 and bcftools call under a haploid model (SNP minimum quality score of 20). Consensus sequences were constructed for each isolate, with positions below 8x depth masked as “N”. Variable sites were extracted using snp-sites v2.5.1 (30) producing an alignment of 55 taxa. IQTREE2 v3.1.2 (31) was used to infer a maximum-likelihood phylogeny under a General Time Reversible model with discrete Gamma distribution and ascertainment bias correction (GTR+ASC+G) and 1000 ultrafast bootstrap replicates, from 49,067 sites (47,270 were parsimony-informative). The phylogeny was midpoint-rooted and visualized with the iTOL online tool (32). To identify clade-specific SNPs, C23- 2 was mapped to clade III reference genome B11221 (GenBank accession GCA_031357565.2) using BWA-MEM v0.7.19 (29). SNPs were identified using SAMtools v1.23 (33), and VarScan v2.4.6 (34), setting the minimum coverage of 10 reads and variant allele frequency to .75.

### Pairwise SNP difference

Pairwise SNP differences among the clinical isolates were calculated using the consensus sequences described above, excluding the clade III isolate. A median of 12,382,376 positions was compared for each isolate pair. Pair-wise distances were visualized as a heatmap using Seaborn v0.13.2 and isolates were hierarchically clustered via average linkage (UPGMA) of the distance matrix using SciPy v1.16.3. To compare genetic distances among isolates from the same patient to those between patients, pairwise differences among serial isolates from each patient were averaged to yield a mean SNP difference per patient and between-patients, diversity was assessed using all pairwise differences among isolates from different patients.

### Growth Kinetics

Isolates were grown overnight in YPD broth at 30°C. Then, 200μl of 1×10^5^ cells/ml in YPD broth was inoculated in wells of 96-well round-bottom microtiter plates and incubated at 37°C for 4h. Absorbance (optical density at 630, OD_630_) was measured using a Cell Imaging Multi-Mode Reader (Cytation 5; Biotek).

### Biofilm formation and aggregation

Biofilm formation was assessed based on metabolic activity as we previously described (26). Briefly, 200µl of 1×10^6^ cells/ml in RPMI1640 medium buffered with HEPES (Difco) were seeded in wells of flat-bottom 96-well polystyrene microtiter plates. Following incubation at 37°C for 24h, wells were washed twice with phosphate-buffered saline (PBS) and biofilms evaluated using the MTS metabolic assay (Promega); color intensity was measured at OD_490_ using a Biomate 3S spectrophotometer (Thermo). Aggregation was assessed based on formation of cell aggregates. Cell suspensions at 5×10^7^ cells/ml density were vigorously vortexed for 1 min and formation of cell aggregates monitored visually.

### RNA extraction

RNA extraction was performed using the RiboPure Yeast Kit (Thermo Scientific) following manufacturer instructions. Briefly, cells were grown overnight at 37°C in YPD broth and harvested by centrifugation. Cell pellets were resuspended in Lysis Buffer with Zirconia Beads, vortexed for 10min and cell lysate clarified by centrifugation to remove cell debris. Eluted RNA was treated with DNase I (8 U) and analyzed in a Nanodrop Lite (Thermo Scientific).

### RT-qPCR

The transcript abundance of *ALS4112* and *SCF1* genes was evaluated for 10 representative isolates categorized as aggregative and robust biofilm formers (Agg+/RBF) vs non-aggregative and poor biofilm formers (Agg-/PBF) (5 isolates in each group). The non-aggregative and poor biofilm former AR0387 strain was used as reference. Extracted RNA was used to synthesize cDNA using High-Capacity cDNA Reverse Transcription Kit (Applied Biosystems) following manufacturer instructions. The generated cDNA served as the template for qPCR reactions using the Power-Up SYBR Green Master Mix (Applied Biosystems). Amplification was carried out using a QuantStudio 3 Real-Time PCR System (Applied Biosystems). Oligonucleotide primers specific to the target genes were designed using NCBI Primer-BLAST with the *C. auris* B8441 genome assembly (NCBI GCA_002759435.2) as a reference. *ACT1* (B9J08_000486), *ALS4112* (B9J08_004112) and *SCF1* (B9J08_001458) genes were amplified using primers listed in Supp Table 1. Experiments were performed on 3 biological replicates with technical triplicate. The 2^−ΔΔ^*CT* method was used to calculate relative levels of expression of each gene normalized to *ACT1* gene. Fold change values are expressed relative to strain AR0387 with error bars representing standard errors of the means. Statistical comparisons were made using Welch’s ANOVA with Dunnett T3 post hoc test.

### Data analysis

Statistical analysis of biofilm growth was performed using R statistical programming software. To compare differences among strains in biofilm forming capabilities and growth rates, a oneway ANOVA with Tukey’s post host test was used. To compare gene expressions, Welch’s ANOVA with Dunnett T3 post hoc test was used due to significant different SDs among groups. To compare genetic diversity within and between patients, groups were compared using Mann-Whitney U test. *P* values less than 0.05 were considered significant. Ggplot2 and ggpubr packages and Matplotlib library were used to construct figures.

## RESULTS

### Antifungal susceptibilities of clinical isolates

All 53 isolates were found to be resistant to fluconazole (100%), six isolates were resistant to amphotericin B (11%), eight isolates (15%) from six unique patients were resistant to caspofungin and micafungin, and four (7.5 %) of the isolates from four unique patients were resistant to all four antifungals tested (Table 1). Among the six patients with resistant isolates, three had been treated with micafungin prior to recovery of the evaluated isolates.

### Whole genome sequencing and phylogenetic analysis

Maximum-likelihood pairwise distances confirmed that C23-2 and South African clade III reference genome B11211 were each other’s nearest neighbors (distance ≈ 0.016) and that the other 52 isolates formed a single clade with the South Asian clade I B8441 reference genome (Fig. 1A).

**Figure 1.**
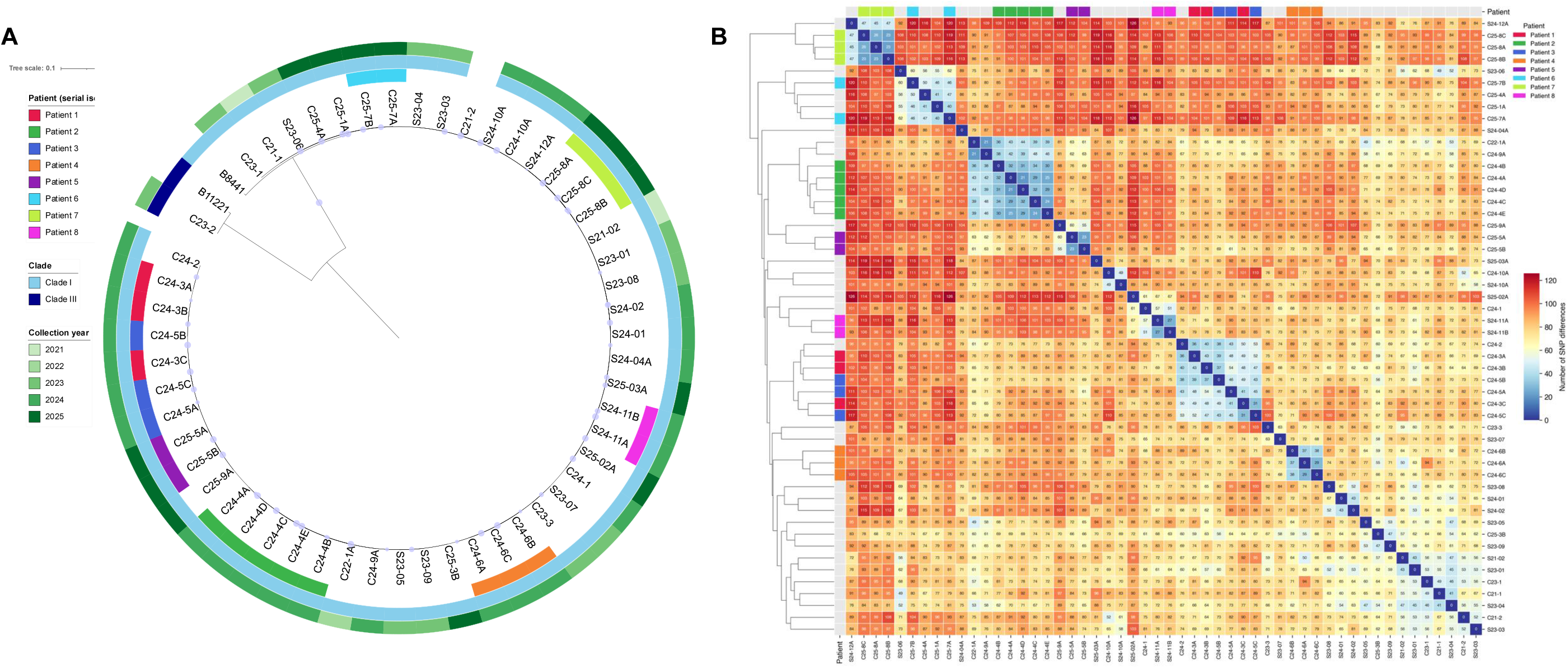
Phylogenetic tree and pairwise SNP relationships among clinical isolates. **(A)** Maximum-likelihood phylogeny of 53 clinical isolates and clade I and clade III representative genomes B8441 and B11221. Isolates are labelled by colored rings with the innermost ring indicating collection year, middle ring indicating patient origin for serial isolates, and outer ring indicating clade. **(B)** Heatmap of hierarchical clustering of 52 clade I isolates based on pairwise genetic distances. Isolates obtained from same patient are color-coded. Scale Bar shows pairwise SNP distances with blue representing smaller genetic distances and red as larger genetic distances.

### Genetic Diversity

From the distance matrix, most serial isolates from patients clustered together, consistent with the maximum-likelihood phylogeny. However, isolates from three different patients (C24-2, C24-3 and C24-5) did not separate into three distinct clusters and two isolates (C25-7A and C25-7B) from the same patient, although genetically close (46 SNP difference), did not form one cluster (Fig. 1B). The median pairwise SNP difference among isolates was 83 SNPs (mean 82.3). Genetic diversity among serially collected isolates from eight patients (mean 33.3 SNPs across 25 pairs) was significantly lower than that among isolates from different patients (mean 83.2 across 1,301 pairs, p ≤.0001).

### Mutations

Whole genome sequencing (Fig. 2) demonstrated that all isolates have the Y132F in *ERG11* and the E709D (35) in *CDR1*. In *TAC1B* gene, all the isolates had the reported A583S mutation, however, 4 of the isolates had an additional T385I substitution in the *TAC1B* gene not previously reported. Additionally, among the 8 isolates resistant to echinocandins, one isolate had the non-synonymous mutation resulting in the F635Y substitution in *FKS1* whereas 4 isolates had a previously reported D642Y substitution (36-38); 3 of these 4 isolates were recovered from the same patient. In one echinocandin resistant isolate (C25-3B), low-frequency D642Y substitution (31%) was observed but was below the calling threshold. In another echinocandin resistant isolate, only one mutation (R641G) not previously associated with resistance was found in *FKS1*. In one resistant isolate, an M690I mutation in hot spot region 3 was the only one found in *FKS1*. The M690I substitution was also found in a susceptible isolate (24-4D) from a patient with multiple isolates recovered (24-4A-E) where the subsequent isolate (24-4E) recovered 8 months later became multi-drug resistant and exhibited the *FKS1* F635Y substitution (Table 1; Fig. 2). No known mutations in studied ergosterol (ERG) genes associated with amphotericin B resistance were identified however, a W96L mutation was found in the *ERG25* gene in 4 of the echinocandin resistant isolates.

**Figure 2.**
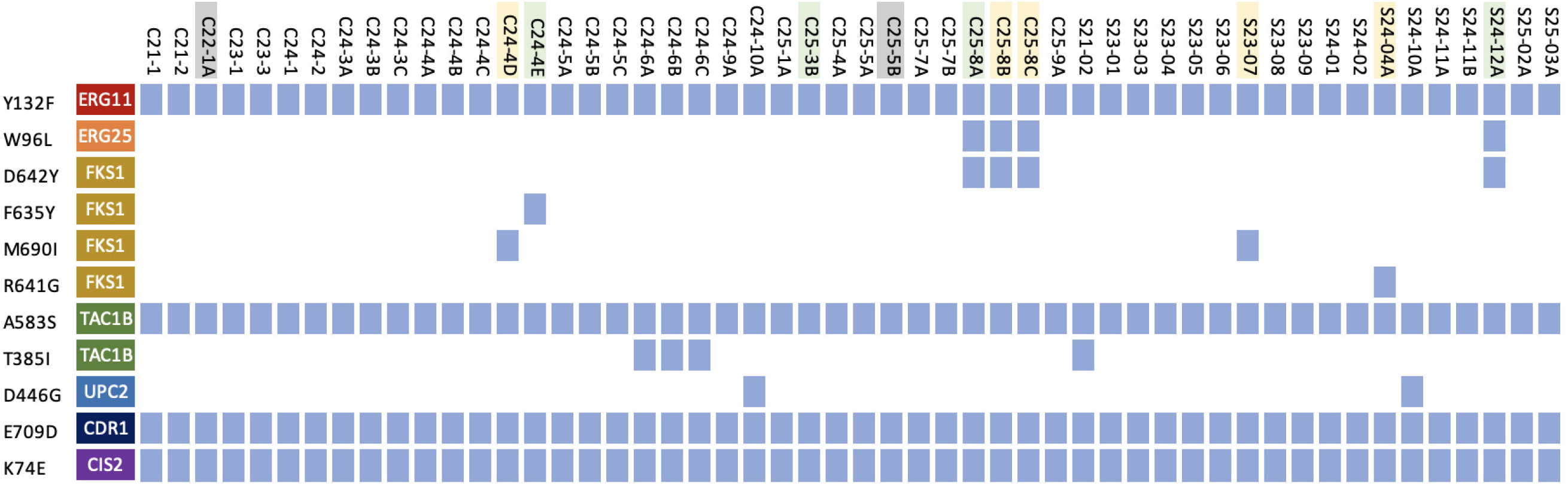
Substitutions in genes associated with antifungal resistance. Whole genome sequencing reveals amino acid substitutions in genes associated with antifungal resistance including those within hotspots 1 and 3 of *FKS1*. All isolates were resistant to fluconazole. Gray isolates: resistant to amphotericin B; yellow isolates resistant to echinocandins; green isolates are multi-drug resistant.

### Isolates vary in growth rates, biofilm forming abilities and aggregation

Isolates exhibited varied growth rates including among some isolates recovered from same patient (Supp. Fig. 2), but no significant associations were seen between growth rates and drug susceptibility or other variables. Similarly, isolates varied widely in their abilities to form biofilms (Fig. 3). Eight patients had multiple isolates recovered; in four of these patients, no differences in biofilm formation were found among each patient’s respective isolates. However, significant differences were seen among isolate sets recovered from four patients: among the five isolates from patient 24-4, only the second and last had significant (p<0.0086) difference in biofilm formation; for patients 24-3 and 24-5, all three isolates exhibited significant differences from each other (p=.041, p=0.00012, p=0.0000248, p=0.018, p=0.0084, p=0.00034, respectively) and for patient 25-8, the first two isolates differed significantly (p = 0.0423 and p=0.11) from the last isolates.

**Figure 3.**
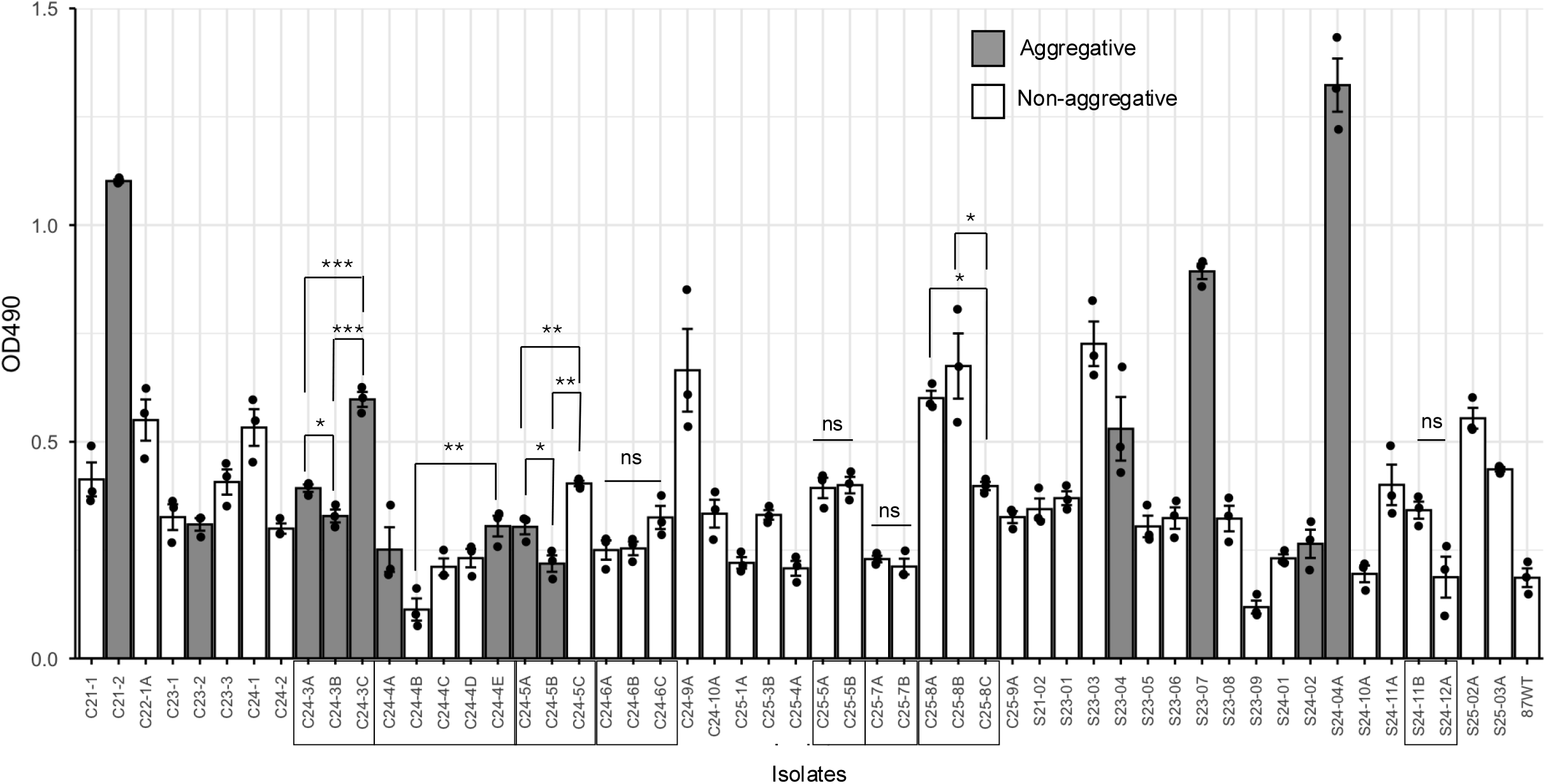
Comparative evaluation of biofilm formation and cell aggregation of clinical isolates. Boxed isolates are recovered from the same patient with statistical differences between the isolates in biofilm formation indicated. Bar-plots show mean and standard error of mean of *n* = 3 biological replicates, each as an average of 3 technical replicates. \**P* ≤ 0.05, \*\**P* ≤ 0.01, \*\*\**P* ≤ 0.001 ns: no significance.

Of the 53 isolates analyzed, 14 demonstrated aggregative properties. The single clade III isolate displayed the highest degree of aggregation, with cell aggregates rapidly forming and sedimenting within 2 mins (Supp. Fig. 3). The 13 clade I isolates exhibited various but markedly lower levels of aggregation to that of the clade III isolate. The aggregative or non-aggregative phenotype were consistent among the isolates within each patient set except in 1 patient (24-4) with 5 isolates where only the first and last isolates aggregated. Similarly, in another patient with 3 isolates, only the first 2 aggregated. No strong associations were found between aggregation and biofilm formation as although all aggregative isolates formed good biofilms, some good biofilm formers were non-aggregative. Similarly, no correlation was found between aggregation and drug resistance (only 2 of the 14 aggregative isolates were resistant) (Table 1).

### Biofilm formation and aggregation is associated with modulation of expression of *SCF1* and *ALS4112* genes

The transcript abundance of the cell wall adhesins genes *ALS4112* and *SCF1* was evaluated between representative isolates categorized as aggregative and robust biofilm formers (Agg+/RBF) vs non-aggregative and poor biofilm formers (Agg-/PBF) using the non-aggregative and poor biofilm former AR0387 strain as the reference. The expression of both *ALS4112* (P < 0.0001) and *SCF1* (P < 0.01) had a strong positive association with aggregation and biofilm formation (Fig. 4A, C). On average, the Agg+/RBF group had >8-fold increase in expression of *ALS4112* and >250-fold increase in *SCF1* genes compared to the Agg-/PBF group (Fig. 4B, D). However, one aggregative and robust biofilm former isolate (C23-07), although exhibited high *ALS4112* expression, *SCF1* expression was comparable to that in the Agg-/PBF isolates (Fig. 4C).

**Figure 4.**
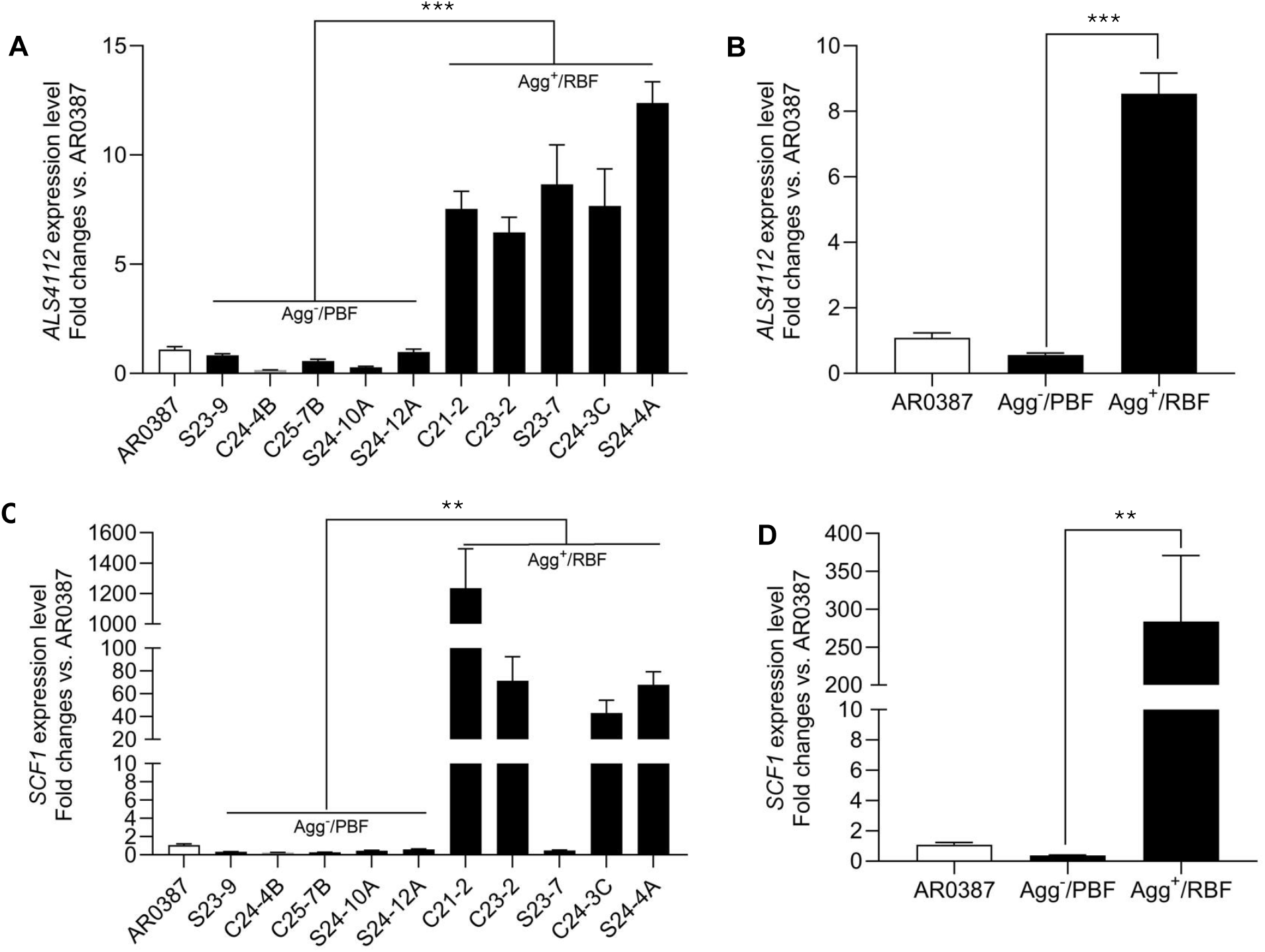
Expression of cell wall adhesin genes in representative clinical isolates. Comparative *ALS4112* **(A, B)** and *SCF1* **(C, D)** gene expression among non-aggregative and poor biofilm forming isolates (Agg^-^/PBF) and aggregative and robust biofilm forming isolates (Agg^+^/RBF). Relative levels of expression of each gene were normalized to *ACT1* gene expression and fold change values are expressed relative to the non-aggregative and poor biofilm forming AR0387 reference strain. Data presented is the average of experiments performed in triplicate on 3 separate occasions. Error bars represent standard errors. \*\**P* ≤ 0.01, \*\*\**P* ≤ 0.001.

## DISCUSSION

Due to its remarkable adaptability, transmissibility, and capacity to persist on biotic and abiotic surfaces *C. auris* has emerged as a global multidrug-resistant threat. In this study, we investigated the genetic diversity and resistance mechanisms of *C. auris* clinical isolates. Importantly, we aimed to provide mechanistic insights into the observed phenotypic plasticity among circulating *C. auris* isolates focusing on aggregation and biofilm formation. Based on phylogenetic analysis, 52 isolates belonged to clade I and only one isolate belonged to clade III. Importantly, seven clade I isolates recovered from three patients (C24-2, C24-3 and C24-5) were found to be genetically closely related and phenotypically similar compared to the other isolates. Interestingly, the three patients were all transplant patients present in the surgical intensive care unit at the same time and passed away within days of each other raising the prospect of patient-to-patient transmission.

Susceptibility testing of *C. auris* isolates demonstrated all isolates to be resistant to fluconazole, consistent with previous observations that South Asia clade I isolates are overwhelmingly resistant to fluconazole (13). Mutations in *ERG11, TAC1B*, and *CDR1* genes have important roles for azole resistance with the most commonly reported mechanism is the acquisition of mutations in *ERG11*, the gene encoding lanosterol 14α-demethylase, and all of our isolates were found to have the Y132F mutation (35, 39). Additionally, isolates also had mutations in the efflux pump gene *CDR1* and the *TAC1B* gene that encodes transcription factors that regulate the expression of efflux pump genes (35, 40, 41). Interestingly, one mutation in the *TAC1B* gene not previously reported (T385I) was found in four of the isolates.

While echinocandin resistance remains uncommon, recent reports demonstrate an increase in such cases. Among our collection, eight (15%) isolates were found to be resistant to echinocandins (caspofungin and micafungin) harboring previously reported mutations (D642Y and F635Y) in the *FKS1* gene encoding 1,3-β-glucan synthase (36-38, 42). However, in one resistant isolate, R641G was the only mutation found in *FKS1;* although not previously associated with echinocandin resistance in *C. auris*, in *Candida* species, substitution in amino acid positions 635-642 was reported to affect drug binding leading to emergence of resistance (43). Therefore, this yet uncharacterized mutation in *C. auris* may be linked to echinocandin resistance. In one echinocandin resistant isolate (C25-3B) a low frequency (31%) of the D642Y substitution was observed although it did not meet our variant-calling threshold. This is interesting as it may indicate the presence of mixed population of resistant and susceptible cells consistent with hetero-resistance, or an emerging resistant subpopulation under echinocandin therapy. In another echinocandin resistant isolate, we found an M690I mutation in hot spot region 3 of the *FKS1* gene. This substitution was previously reported by Carolus et al. (44) during in vitro evolution experiments of acquired resistance to caspofungin and more recently in resistant and susceptible isolates by other studies (36, 45). In developing a benchmark dataset of sequenced *C. auris* isolates to detect echinocandin resistance-conferring mutations in *FKS1*, Misas et al. (36) found resistant isolates with no known *FKS1* mutations harboring the M690I amino acid change, which they also found in a susceptible isolate. Similar to these findings, in addition to the resistant isolate, we also found this M690I mutation in a susceptible isolate. Interestingly however, although susceptible, the subsequent isolate recovered from the same patient who was receiving antifungal therapy, developed multi-drug resistance with an F635Y mutation in the *FKS1* gene. These findings underscore the threat of therapy-induced development of resistance and warrant further studies into the potential contribution of the M690I variant to the development of echinocandin resistance. An additional noteworthy observation was the presence of a W96L mutation in the *ERG25* gene in 5 of the echinocandin resistant isolates harboring the D642Y mutation in the *FKS1*. Although the relevance is not clear, in *C. albicans, ERG251* and *ERG25* are two paralogs with shared activity that encode a predicted C4-sterol methyl oxidase involved in biofilm formation (46).

Similar to echinocandins, resistance to amphotericin B was reported to develop in patients on therapy (19). However, the mechanism for amphotericin B resistance is unclear as only one study identified a mutation in the *ERG6* gene in a clinical isolate to be associated with increased resistance to amphotericin B (19). Among our isolates, only six were resistant to amphotericin B and in line with findings from other studies, we did not identify non-synonymous variants among amphotericin B-susceptible and resistant isolates in studied ergosterol (ERG) genes. Although it is unknown what genetic elements may be driving amphotericin B resistance, reduced membrane lipid permeability or overexpression of ERG genes may have been contributors (3). In terms of multi-drug resistance, 4 of our isolates recovered from four unique patients were found to be resistant to all three classes of antifungals. Most importantly is the demonstration of multi-drug resistance development in a patient undergoing antifungal treatment, underscoring the threat of evolution of drug resistance induced by antifungal therapy.

Phenotypic evaluation of the isolates indicated wide variations in biofilm formation and aggregation including among some isolates recovered from the same patients. It is reported that *C. auris* strains aggregate in vitro by exposure to antifungals, but this form of aggregation is consistent with a cell separation defect due to cell wall changes (47). Therefore, it is possible that aggregate formation may constitute a protective response induced by therapy to impede drug penetration into the cell aggregates. However, another form of aggregation is induced by changes in expression of genes associated with adhesion (26, 47-49). We recently identified two cell wall adhesins, Scf1 and Als4112 as key mediators for biofilm formation and aggregation with Als4112 being more crucial for aggregation (26).

To provide some mechanistic insights behind the diversity of the clinical isolates in their abilities to aggregate and adhere, we investigated the gene expression of these two key adhesins in representative isolates exhibiting the various phenotypes. As expected, findings strongly associated *ALS4112* and *SCF1* expression with aggregation and biofilm formation indicating that variability among *C. auris* clinical isolates in these phenotypic characteristics is primarily mediated by differential expression of cell wall adhesins. However, one aggregative and robust biofilm forming isolate exhibited high expression of *ALS4112* but not *SCF1*. These findings are interesting as we have demonstrated complementary roles for these adhesins, which despite their limited similarity, may function redundantly to promote cell-cell interaction and biofilm formation (26). Therefore, the abundance of Als4112 in that isolate could have compensated for low Scf1 levels. This functional diversity of cell wall adhesins first demonstrated here among clinical isolates, may provide *C. auris* with flexibility and rapid adaptation to the environment.

The aggregation phenotype has been associated with conferring an advantage for skin colonization in line with findings from a recent study by Zhao et al (50) showing that Als4112 regulates *C. auris* adherence to keratinocytes and facilitates skin colonization. These findings are interesting since aggregation is a phenotype primarily of clade III strains, consistent with our findings as the most highly aggregative isolate among our collection with high *ALS4112* expression was the single clade III isolate. Further, our clade III isolate was recovered from infected tissue indicating that skin colonization may have led to infection. This morphological plasticity likely represents a strategy that allows this pathogen to adapt to changing environmental conditions and survive in certain host niches.

Despite major advances in our knowledge of *C. auris* biology and resistance mechanisms, significant knowledge gaps remain in our understanding of how *C. auris* adapts to environmental pressures and evolves drug resistance. With *C. auris* infections and incidence of drug resistance concerningly on the rise, understanding these adaptive characteristics is key to developing effective clinical and therapeutic management strategies.

## ACKNOWLEDGEMENTS

The work in this publication was supported by the University of Maryland Baltimore, Institute for Clinical & Translational Research (ICTR) and in part by 1R25AI181751-01 from the NIH National Institute of Allergy and Infectious Diseases. The funders had no role in study design, data collection and interpretation, or the decision to submit the work for publication. We would like to thank Harris Levy for his assistance.

## DATA AVAILABILITY

Upon acceptance and prior to publication, all of the raw sequencing reads from this study will be available at the NCBI Sequence Read Archive (SRA) under BioProject accession number PRJNA1086003.

## SUPPLEMENTAL MATERIAL

Supp table 1. Primers used in this study.

**Supp Figure 1. Pairwise SNP differences within and between patients. (A)** Mean pairwise single nucleotide polymorphism difference among serially collected isolates (patients with at least 2 isolates recovered) and calculated as the mean of all pairwise comparisons within that set (n=8). **(B)** Individual pairwise single nucleotide differences within (n=25) and among patients (n=1301). Median and interquartile range are depicted here by horizontal bars. Statistical comparison was made using Mann-Whitney U test (*P* ≤ 0.001***).

**Supp Figure 2. Growth kinetics**. Evaluation of cell growth rates among isolates over 36h based on values of OD_630_. Percent change in growth rate from 0 to 4 hours of growth are shown. Boxed isolates are recovered from the same patient with statistical differences in growth rate among the isolates indicated. Statistical analysis was performed by one-way ANOVA and post-hoc Tukey test. Bar-plots show mean and standard error of mean of *n* = 3 biological replicates, each as an average of 3 technical replicates. \**P* ≤ 0.05, \*\**P* ≤ 0.01, \*\*\**P* ≤ 0.001 ns: no significance.

**Supp Figure 3. Cell aggregation**. Visible formation of cell aggregates and rapid sedimentation of clade III aggregative (Agg+) isolate (C23-2) compared to a non-aggregative (Agg-) isolate.

## REFERENCES

1. Forsberg K, Woodworth K, Walters M, Berkow EL, Jackson B, Chiller T, Vallabhaneni S. 2019 Candida auris: The recent emergence of a multidrug-resistant fungal pathogen. Med Mycol 57:1–12. .

2. Lockhart SR, Etienne KA, Vallabhaneni S, Farooqi J, Chowdhary A, Govender NP, Colombo AL, Calvo B, Cuomo CA, Desjardins CA, Berkow EL, Castanheira M, Magobo RE, Jabeen K, Asghar RJ, Meis JF, Jackson B, Chiller T, Litvintseva AP. 2017. Simultaneous emergence of multidrug-resistant Candida auris on 3 continents confirmed by whole-genome sequencing and epidemiological analyses. Clin Infect Dis 64:134–140.

3. Chowdhary A, Jain K, Chauhan N. 2023. Candida auris genetics and emergence. Annu Rev Microbiol 77:583–602.

4. Lionakis MS, Chowdhary A. 2024. Candida auris infections. New Eng J Med 39:1924–1935.

5. Lyman M, Forsberg K, Sexton DJ, Chow NA, Lockhart SR, Jackson BR, Chiller T. 2023. Worsening Spread of Candida auris in the United States, 2019 to 2021. Ann Intern Med 176:489–495.

6. Vaccaro A, Cooper JF, Vazquez-Rodriguez A, Badali H, Kean R, Ramage G, Lopez-Ribot JL. 2026 Candidozyma auris and the perfect storm of fungal pathogenicity: Adaptation, persistence, and resistance. J Fungi (Basel) 12.

7. Chow NA, Muñoz JF, Gade L, Berkow EL, Li X, Welsh RM, Forsberg K, Lockhart SR, Adam R, Alanio A, Alastruey-Izquierdo A, Althawadi S, Araúz AB, Ben-Ami R, Bharat A, Calvo B, Desnos-Ollivier M, Escandón P, Gardam D, Gunturu R, Heath CH, Kurzai O, Martin R, Litvintseva AP, Cuomo CA. 2020. Tracing the evolutionary history and global expansion of Candida auris using population genomic analyses. mBio 11:e03364–19.

8. Thangamani S, Balakumar A, Datta A, Bryak G, Lionakis MS. 2026 Host-Candida auris interactions in the skin. PLoS Pathog 22:e1014075.

9. Kean R, Brown J, Gulmez D, Ware A, Ramage G. 2020. Candida auris: A decade of understanding of an enigmatic pathogenic yeast. J Fungi (Basel) 6:30.

10. de Jong AW, Hagen F. 2019. Attack, defend and persist: How the fungal pathogen Candida auris was able to emerge globally in healthcare environments. Mycopathologia 148:353–365.

11. Welsh RM, Bentz ML, Shams A, Houston H, Lyons A, Rose LJ, Litvintseva AP. 2017. Survival, persistence, and isolation of the emerging multidrug-resistant pathogenic yeast Candida auris on a plastic health care surface. J Clin Microbiol 55:2996–3005.

12. Organization WH. 2022. WHO fungal priority pathogens list to guide research, development and public health action. Geneva: WHO.

13. Rhodes J, Jacobs J, Dennis EK, Manjari S R., Banavali NK, Marlow R, Rokebul MA, Chaturvedi S, Chaturvedi V. 2024 What makes Candida auris pan-drug resistant? Integrative insights from genomic, transcriptomic, and phenomic analysis of clinical strains resistant to all four major classes of antifungal drugs. Antimicrob Agents Chemother 68:e0091124.

14. Wang WW, Putnam NE, Johnson JK, Jabra-Rizk MA. 2025. In vivo evolution of Candida auris multi-drug resistance in a patient receiving antifungal treatment. J Infect Dis 232:1203–1207.

15. Tian S, Wu Y, Li H, Rong C, Wu N, Chu Y, Jiang N, Zhang J, Shang H. 2024. Evolutionary accumulation of FKS1 mutations from clinical echinocandin-resistant Candida auris. Emerg Microbes Infect 13:2377584.

16. Spruijtenburg B, Ahmad S, Asadzadeh M, Alfouzan W, Al-Obaid I, Mokaddas E, Meijer EFJ, Meis JF, de Groot T. 2023 Whole genome sequencing analysis demonstrates therapy-induced echinocandin resistance in Candida auris isolates. Mycoses 66:1079–1086.

17. Jacobs SE, Jacobs JL, Dennis EK, Taimur S, Rana M, Patel D, Gitman M, Patel G, Schaefer S, Iyer K, Moon J, Adams V, Lerner P, Walsh TJ, Zhu Y, Anower MR, Vaidya MM, Chaturvedi S, Chaturvedi V. 2022 Candida auris pan-drug-resistant to four classes of antifungal agents. Antimicrob Agents Chemother 66:e0005322.

18. Shivarathri R, Jenull S, Chauhan M, Singh A, Mazumdar R, Chowdhary A, Kuchler K, Chauhan N. 2022. Comparative transcriptomics reveal possible mechanisms of amphotericin B resistance in Candida auris. Antimicrob Agents Chemother 66:e0227621.

19. Rybak JM, Barker KS, Muñoz JF, Parker JE, Ahmad S, Mokaddas E, Abdullah A, Elhagracy RS, Kelly SL, Cuomo CA, Rogers PD. 2022. In vivo emergence of high-level resistance during treatment reveals the first identified mechanism of amphotericin B resistance in Candida auris. Clin Microbiol Infect 28:838–843.

20. Lockhart SR. 2019 Candida auris and multidrug resistance: Defining the new normal. Fungal Genet Biol 131:103243.

21. Chaabane F, Graf A, Jequier L, Coste AT. 2019. Review on antifungal resistance mechanisms in the emerging pathogen Candida auris. Front Microbiol 10:2788.

22. Kean R G. R. 2019. Combined antifungal resistance and biofilm tolerance: the Global threat of Candida auris. mSphere 4.

23. Vila T, Montelongo-Jauregui D, Hussian A, Puthran T, Sultan AS, Jabra-Rizk MA. 2020. Comparative evaluations of the pathogenesis of Candida auris phenotypes and Candida albicans using clinically relevant murine models of infections. mSphere 5:e00760–20.

24. Brown JL, Delaney C, Short B, Butcher MC, McKloud E, Williams C, Kean R, Ramage G. 2020 Candida auris phenotypic heterogeneity determines pathogenicity in vitro mSphere 5:e00371–20.

25. Pelletier C, Shaw S, Alsayegh S, Brown AJP, Lorenz A. 2024 Candida auris undergoes adhesin-dependent and -independent cellular aggregation. PLoS Pathog 20:e1012076.

26. Wang TW, Sofras D, Montelongo-Jauregui D, Paiva TO, Carolus H, Dufrêne YF, Alfaifi AA, McCracken C, Bruno VM, Van Dijck P, Jabra-Rizk MA. 2024. Functional Redundancy in Candida auris cell surface adhesins crucial for cell-cell interaction and aggregation. Nat Commun 15:9212.

27. Forgács L, Borman AM, E. p, al. e. 2020. Comparison of in vivo pathogenicity of four Candida auris clades in a neutropenic bloodstream infection murine model. Emerg Microbes Infect 9:1160–1169.

28. Bing J, Guan Z, Zheng T, Ennis CL, Nobile CJ, Chen C, Chu H, Huang G. 2024 Rapid evolution of an adaptive multicellular morphology of Candida auris during systemic infection. Nat Commun 15:2381.

29. Li H, Durbin R. 2009. Fast and accurate short read alignment with Burrows-Wheeler transform. Bioinformatics 25:1754–60.

30. Page AJ, Taylor B, Delaney AJ, Soares J, Seemann T, Keane JA, Harris SR. 2016. SNP-sites: rapid efficient extraction of SNPs from multi-FASTA alignments. Microbial Genomics 2:e000056.

31. Nguyen LT, Schmidt HA, Von Haeseler A, Minh BQ. 2015. IQ-TREE: a fast and effective stochastic algorithm for estimating maximum-likelihood phylogenies. Molec Biol Evolution 32:268–274.

32. Letunic I, Bork P. 2021. Interactive Tree of Life (iTOL) v5: an online tool for phylogenetic tree display and annotation. Nucleic Acids Research 49:W293–W296.

33. Li H, Handsaker B, Wysoker A, Fennell T, Ruan J, Homer N, Marth G, Abecasis G, Durbin R, Subgroup GPDP. 2009. The sequence alignment/map format and SAMtools. Bioinformatics 25:2078–2079.

34. Koboldt DC, Chen K, Wylie T, Larson DE, McLellan MD, Mardis ER, Weinstock GM, Wilson RK, Ding L. 2009. VarScan: variant detection in massively parallel sequencing of individual and pooled samples. Bioinformatics 25:2283–2285.

35. Esquivel BD, Santos A, Rybak JM, Santana DJ, Rogers PD, White TC. 2026 Mutations in ERG11, TAC1B, and CDR1 reduce fluconazole accumulation in drug-resistant Candidozyma auris isolates. mBio 17:e0395725.

36. Misas E, Parnell LA, Rajeev M, López LF, Santos AR, Mudge ZB, Gade L, Forsberg K, Lyman M, Sexton DJ, Litvintseva AP, Lockhart SR, Chow NA. 2025. A benchmark dataset for validating FKS1 mutations in Candida auris. Microbiol Spectr 13:e0314724.

37. Hirayama T T. M, Sumiyoshim M, Ito Y, Ashizawa N, Takeda K, al. e. 2023. Echinocandin resistance in Candida auris occurs in the murine gastrointestinal tract due to FKS1 mutations. Antimicrob Agents Chemother 67:e01243–22.

38. Asadzadeh M, Mokaddas E, Ahmad S, Abdullah AA, de Groot T, Meis JF, Shetty SA. 2022. Molecular characterisation of Candida auris isolates from immunocompromised patients in a tertiary-care hospital in Kuwait reveals a novel mutation in FKS1 conferring reduced susceptibility to echinocandins. Mycoses 65:331–343.

39. Rybak JM, Sharma C, Doorley LA, Barker KS, Palmer GE, Rogers PD. 2021 Delineation of the direct contribution of Candida auris ERG11 mutations to clinical triazole resistance. Microbiol Spectr 9:e0158521.

40. Rybak JM, Cuomo CA, Rogers PD. 2022. The molecular and genetic basis of antifungal resistance in the emerging fungal pathogen Candida auris. Curr Opin Microbiol 70.

41. Chen XF, Zhang H, Liu LL, Guo LN, Liu WJ, Liu YL, Li DD, Zhao Y, Zhu RY, Li Y, Dai RC, Yu SY, Li J, Wang T, Dou HT, Xu YC. 2024. Genome-wide analysis of in vivo-evolved Candida auris reveals multidrug-resistance mechanisms. Mycopathologia 189.

42. Sharma D, Paul RA, Rudramurthy SM, Kashyap N, Bhattacharya S, Soman R, Shankarnarayan SA, Chavan D, Singh S, Das P, Kaur H, Ghosh AK, Prasad R, Sanyal K, Chakrabarti A. 2022. Impact of FKS1 genotype on echinocandin in vitro susceptibility in Candida auris and in vivo response in a murine model of infection. Antimicrob Agents Chemother 66:e0165221.

43. Gifford H, al e. 2026. Global genomic epidemiology of Candida auris : analysis of 12,644 whole genome sequences from 1997-2024. bioRxiv 20260203703534; doi: https://doiorg/1064898/20260203703534.

44. Carolus H, Pierson S, Muñoz J, Subotić A, Cruz RB, Cuomo CC, Van Dijck P. 2021. Genome wide analysis of experimentally evolved Candida auris reveals multiple novel mechanisms of multidrug resistance. mBio 12:e03333–20.

45. Smithgall MC, Kilic A, Weidmann M, Ofori K, Gu Y, Koganti L, Mi S, Xia H, Shi J, Pang J, Mansukhani M, Hsiao S, Wu F. 2025. Genetic and phenotypic intra-clade variation in Candida auris isolated from critically ill patients in a New York City tertiary care center. Clin Chem 71:185–191.

46. Xiong L, Pereira De Sa N, Zarnowski R, Huang MY, Mota Fernandes C, Lanni F, Andes DR, Del Poeta M, Mitchell AP. 2024 Biofilm-associated metabolism via ERG251 in Candida albicans. PLoS Pathog 20:e1012225.

47. Szekely A, Borman AM, Johnsona EM. 2019. Candida auris isolates of the southern asian and south african lineages exhibit different phenotypic and antifungal susceptibility profiles in vitro. J Clin Microbiol 57:1–12.

48. Santana DJ, Anku JAE, Zhao G, Zarnowski R, Johnson CJ, Hautau H, Visser ND, Ibrahim AS, Andes D, Nett JE, Singh S, O’Meara TR. 2023. A Candida auris-specific adhesin, Scf1, governs surface association, colonization, and virulence. Science 381:1461–1467.

49. Bing J, Guan Z, Zheng T, Zhang Z, Fan S, Ennis CL, Nobile CJ, Huang G. 2023. Clinical isolates of Candida auris with enhanced adherence and biofilm formation due to genomic amplification of ALS4. PLoS Pathog 19:e1011239.

50. Zhao G, Lyu J, Veniaminova NA, Zarnowski R, Mattos E, Johnson CJ, Quintanilla D, Hautau H, Hold LA, Xu B, Anku JAE, Dasgupta K, Hale JJ, Steltzer SS, Santana DJ, Ibrahim AS, Snitkin ES, Andes D, Nett JE, Singh S, Abraham AC, Killian ML, Kahlenberg JM, Wong SY, O’Meara TR. 2025. Adhesin Als4112 promotes Candida auris skin colonization through interactions with keratinocytes and extracellular matrix proteins. Nat Commun 16:5673.

